# The role of aneuploidy and polyclonality in the adaptation of the Protozoan parasite *Leishmania* to high drug pressure

**DOI:** 10.1101/2022.12.22.521483

**Authors:** Gabriel H. Negreira, Robin de Groote, Dorien Van Giel, Pieter Monsieurs, Ilse Maes, Geraldine de Muylder, Frederik Van den Broeck, Jean-Claude Dujardin, Malgorzata A. Domagalska

## Abstract

Aneuploidy is generally considered harmful, but in some microorganisms, it can act as an adaptive mechanism against environmental stresses. Here, we used *Leishmania* – a protozoan parasite with a remarkable genome plasticity – to study the early evolution of aneuploidy under high drug pressure (antimony or miltefosine) as stressor model. By combining single-cell genomics, lineage tracing with cellular barcodes and longitudinal genome characterization, we revealed that antimony-induced aneuploidy changes result from the polyclonal selection of pre-existing karyotypes, complemented by further and rapid de novo alterations in chromosome copy number along evolution. In the case of miltefosine, early parasite adaptation was associated with independent pre-existing point mutations in a miltefosine transporter gene and aneuploidy changes only emerged later, upon exposure to increased concentration of the drug. Thus, polyclonality and genome plasticity are hallmarks of parasite adaptation, but the scenario of aneuploidy dynamics is dependent on the nature and strength of the environmental stress as well as on the existence of other pre-adaptive mechanisms.

## Introduction

Euploidy is the standard genome configuration in eukaryotes, with most genomes containing two homologous sets for each chromosome. Conversely, aneuploidy, i.e., a dosage imbalance between chromosomes in a cell, is commonly lethal or associated with deleterious effects, in particular in multicellular organisms (1, 2). In some unicellular eukaryotes however, aneuploidy can be well-tolerated or even beneficial. It can be found in pathogenic and non-pathogenic unicellular eukaryotes, including *Saccharomyces cerevisiae, Candida albicans, Cryptococcus neoformans* and *Giardia intestinalis* (3–5), with specific aneuploidy changes being able to confer resistance against environmental stresses such as drug pressure. For instance, in *C. albicans*, aneuploidy affecting chromosome 2 promotes cross-resistance against hydroxyurea and caspofungin (6). Additionally, aneuploidy is also a hallmark of cancer, where it is also associated with the emergence of therapeutic resistance, either by promoting dosage changes of key genes or by causing delays in drug-targeted cell cycle stages (7, 8).

In recent years, *Leishmania spp*. emerged as a new and unique model for studying aneuploidy and its adaptive role. These protozoan parasites are the causative agents of a group of diseases known as leishmaniases and display a digenetic life cycle characterized by two main forms: the extracellular promastigote, which lives in the midgut lumen of female sand flies, and the amastigote form, which resides inside macrophages and other phagocytic cells in their vertebrate hosts (9). *Leishmania spp*. belongs to one of the earliest diverging branches in the Eukaryota domain (Discoba), and as such, they display several idiosyncratic genomic and molecular features compared to higher eukaryotes. Their genomes lack gene specific RNApol II promoters and are organized in divergent/convergent long polycistronic arrays encompassing hundreds of genes which are not functionally related (10). Transcription is thus initiated at defined chromosomal locations known as strand switch regions, which flank the polycistrons. In this context, gene dosage has a nearly one-to-one impact on transcription (11) and directly affect gene expression, which otherwise is mainly controlled post-transcriptionally.

Unlike the abovementioned organisms where euploid genomes are common, all *Leishmania* genomes analyzed hitherto are aneuploid, with the most basic profile being characterized by a polysomy (usually tetrasomy) in chromosome 31, contrasting with a disomy in the other 33-35 chromosomes (12). Additional dosage changes affecting multiple chromosomes (up to 22 out of 36 chromosomes) are commonly observed in cultured promastigotes (13) and are associated with a fitness gain in vitro (14, 15), but they also occur -to a lower extent-in amastigotes, in vivo (16). Recently, a multi-omics study demonstrated a proportional impact of polysomies on the average expression of proteins encoded by the respective polysomic chromosomes, although (i) some of these proteins showed reduced dosage effect and (ii) several proteins derived from disomic chromosomes were upregulated. Altogether, these protein changes ultimately correlated with metabolic adaptations (17). Moreover, aneuploidy in *Leishmania* is highly dynamic and changes in response to new environments, such as drug pressure, vertebrate host or vector (11, 18). Interestingly, spontaneous karyotypic modifications constantly occur even among sister cells in clonal populations (with frequencies of somy changes estimated in the absence of drug pressure at 0.002-0.003/generation/sequenced cell), a phenomenon known as mosaic aneuploidy (19, 20). It is postulated that mosaic aneuploidy in *Leishmania* generates phenotypic heterogeneity that can serve as substrate for natural selection, facilitating adaptation to different environmental pressures (21), but this remains an open question. Moreover, the clonal dynamics of populations facing strong environmental stresses and its relationship with aneuploidy modifications is currently unknown.

In the present study we aimed to address these questions using a reproducible in vitro evolutionary model to study aneuploidy modulations and karyotype evolution in the context of adaptation to environmental stresses, invoked here by the direct exposure to high concentrations of 2 drugs, trivalent antimonial (Sb^III^) or miltefosine (further called ‘flash selection’). By combining clonal lineage tracing with cellular barcodes and a longitudinal genomic characterization, we revealed that changes in aneuploidy under Sb^III^ pressure have a polyclonal origin, arising from the reproducible survival of a specific set of lineages, which further expand stochastically. Additionally, using single-cell genome sequencing, we could uncover the evolutionary paths that might have led to the emergence of such aneuploidy changes, which involved the selection of pre-existing karyotypes, complemented by further de novo alterations in chromosome copy number along evolution. In contrast, flash selection with miltefosine did not initially change aneuploidy despite promoting a stronger bottleneck compared to antimony, and adaptation was initially driven by different pre-existing missense mutations in a miltefosine transporter gene (LdMT). Aneuploidy modifications only happened after the already-selected populations were exposed to a 4 times higher dosage. Thus, polyclonality and genome plasticity are hallmarks of parasite adaptation, but the scenario of aneuploidy dynamics is dependent on the nature and strength of the environmental stress as well as on the existence of other pre-adaptive mechanisms.

## Results

Here, our main models consisted in directly exposing a *L. donovani* promastigote clonal strain (BPK282) to a very high concentration of Sb^III^ (382 µM) or miltefosine (25 µM to 100 µM) in vitro. For each model – which we refer as ‘flash selection’ – we followed two molecular approaches. On one hand, we characterized parasite molecular adaptations along selection through bulk and single-cell genome sequencing, with a special focus on aneuploidy. Complementarily, we developed a novel approach for tracing lineages of *Leishmania* promastigotes in vitro using cellular barcodes. With cellular barcodes, individual cells are ‘tagged’ with a short, random nucleotide sequence in their genomes which is transferred to daughter cells along cell division. Thus, the progeny of each barcoded cell shares the same barcode sequence, constituting a clonal lineage (further referred simply as ‘lineage’) that can be quantified with amplicon sequencing (Figure S1). We used these two approaches in longitudinal assays along adaptation to our flash selection models to track the evolutionary dynamics of hundreds of lineages under drug pressure, revealing the bottlenecks associated with each model and the relationship between selection of lineages and the emergence of genomic adaptations.

### Flash selection with Sb^III^ leads to rapid changes in aneuploidy

The first flash selection protocol we used was previously developed in the context of a study on resistance to trivalent antimonial (Sb^III^), in which resistant parasites with fully restored growth to wild-type levels were observed after 5 weeks (and passaged every 7 days) of direct exposure to 382 µM Sb^III^ (18). We reproduced the experiment with a barcoded BPK282 and determined by bulk genome sequencing the genomic alterations encountered in the populations along 5 passages under Sb^III^ pressure and in 4 replicates (further called SePOP1-4). The search for SNPs and indels in coding regions did not reveal any consistent change associated with the Sb^III^ pressure compared to the control populations (cPOP1-4) which were maintained with PBS (fig. S2A). We also evaluated intra-chromosomal copy number variations with a specific attention for MRPA genes which encode an ABC transporter involved in Sb^III^ sequestration that are often amplified in Sb^III^ resistant *Leishmania* (18). The BPK282 strain already contains a natural amplification of MRPA genes that may bring a pre-adaptation to Sb^III^ (13) and the locus might be subject to further expansion or contraction. In 3 out of 4 replicates, the copy number of the MRPA locus remained stable at ∼3 copies per haploid genome similarly to the initial condition as well as to cPOP1-4, with the exception being SePOP1, which displayed an increase to almost 10 copies per haploid genome at passage 5 (fig. S2B). When investigating aneuploidy variation, we found that BPK282 contained six chromosomes with somy higher than 2 (chr 5, 9, 16, 23, 26 and 31) at the onset of the experiment, and we found a further dosage increase affecting 5-8 chromosomes at passages 2-3 in SePOP1-4 after Sb^III^ exposure (Fig.1a). Interestingly, although different aneuploidy profiles emerged in different replicates, all 4 replicates consistently shared a dosage increase of chromosomes 23, 27 and 31, which could point to an adaptive advantage of the amplification of these chromosomes.

**Fig. 1.**
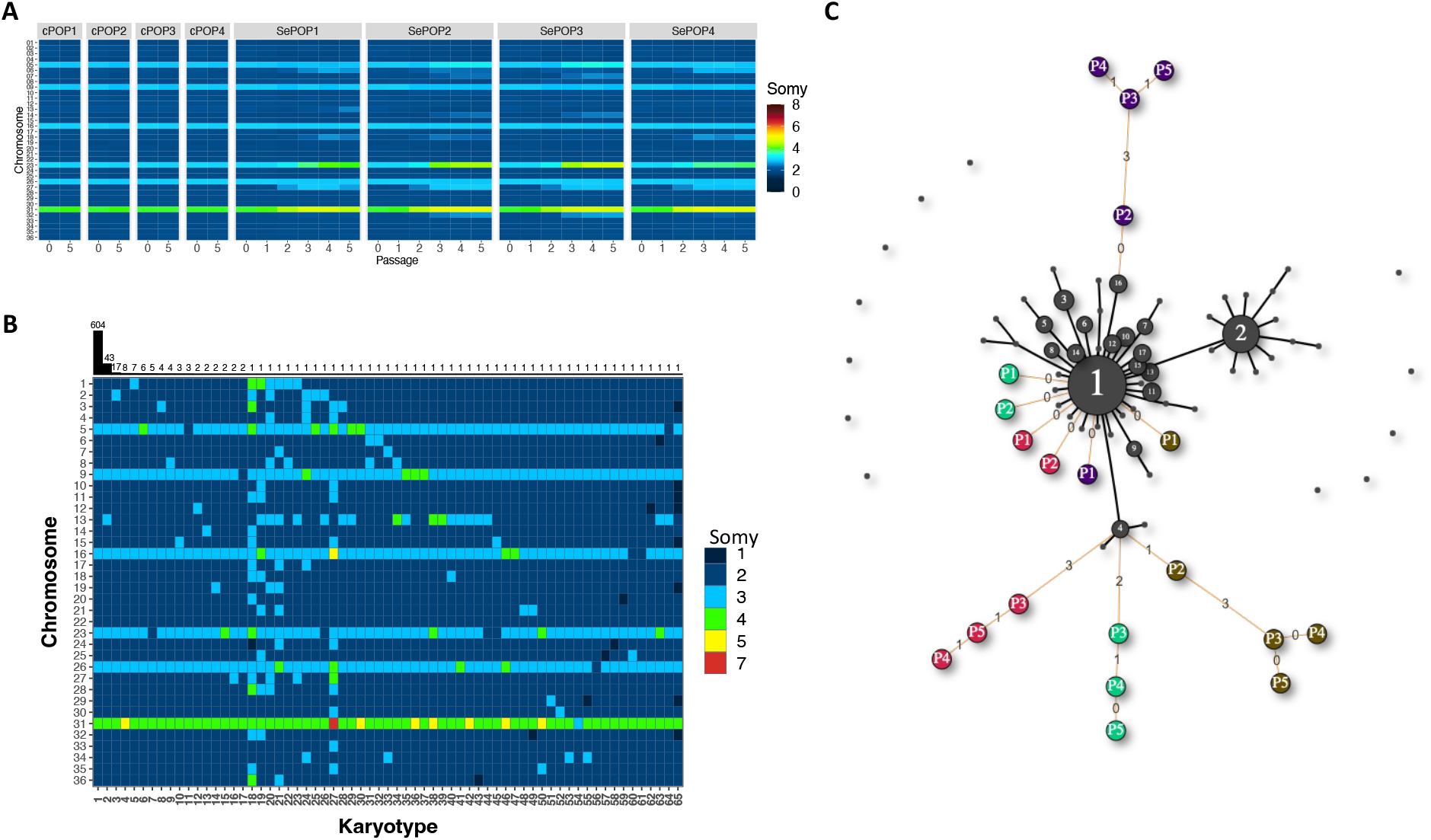
Aneuploidy changes of *L. donovani* BPK282, during flash selection with Sb^III^. **A**. Bulk genome sequencing: Heatmap showing the average copy number of each chromosome between passages 0 (before drug exposure) and passage 5 in cPOP1-4 and along all 5 passages in SePOP1-4. **B**. Heat map depicting all karyotypes identified in the barcoded population prior to Sb^III^-exposure using single-cell genome sequencing. Karyotypes are ordered decreasingly based on their frequency. Bars on the top display the number of promastigotes found with each karyotype. **C**. Minimum spanning tree displaying the number of somy changes between the karyotypes identified in the single-cell data (black nodes) and the rounded bulk aneuploidy of the Sb^III^-selected populations (colored nodes: purple = SePOP1, green = SePOP2, red = SePOP3 and brown = SePOP4) at passages 1-5 (P1-P5). Black lines connecting two nodes indicate that these two karyotypes are different by a single somy change. Orange lines connect the bulk karyotypes of the SePOP1-4 to the single-cell data. Numbers in the orange lines indicate how many somy changes are between the nodes connected by them. Unconnected black nodes are single-cell karyotypes that have 2 or more somy differences compared to any other karyotype.

### Single-cell genomics reveal potential evolutionary paths that led to SbIII-associated aneuploidy changes

To evaluate if the aneuploidy changes observed in the Sb^III^-exposed populations are due to the selection of pre-existing or de novo generated karyotypes, we submitted the same barcoded cell population to high throughput single-cell genome sequencing prior to flash selection. In total, 864 promastigotes were individually sequenced, with a total of 65 different karyotypes (kar1-65) being detected (fig. 1B). These single-cell data revealed a relatively reduced mosaicism, with almost 70% of the parasites displaying the same karyotype (kar1). None of the pre-existing karyotypes were identical to the aneuploidy profiles observed in bulk in SePOP1-4 (fig. 1B). However, individual somy changes consistently observed under Sb^III^ pressure in SePOP1-4 (chromosome 23, 27 and 31) were already present – in few cells – before the flash selection. For instance, kar18 – one of the most aneuploid karyotypes, had a tetrasomic chromosome 23 and a trisomic chromosome 27, while kar38 and kar50 both shared amplification of chromosome 23 and 31. Other single-cells showed dosage increase of one of these chromosomes only, for instance kar15 that only showed tetrasomy of chromosome 23. However, none of the sequenced promastigotes showed amplification of chromosomes 23, 27 and 31 concomitantly, and no pre-existing karyotype was identified with a pentasomy in chromosome 23 as observed in the SePOP3, suggesting that some of these aneuploidy modifications were generated along adaptation to Sb^III^.

To gain insights on possible evolutionary paths that might have led to the emergence of the aneuploidy changes observed in the SePOP1-4 (bulk data) from the initial population (single-cell data), we built a minimum spanning tree connecting the karyotypes found in the single-cell data (fig. 1C – black nodes) to the rounded aneuploidy profiles of SePOP1-4 (fig. 1C – coloured nodes). We based this analysis on the previous observation that the rounded bulk aneuploidy profile of a given promastigote population reflects the most dominant karyotype in that population (20). In this tree, edges connecting nodes represent the number of somy differences between the two connected karyotypes. This approach revealed that the shortest path between pre-existing karyotypes and the selected karyotypes in SePOP1 (purple nodes) starts from kar16, which has exactly the same aneuploidy profile as the one observed in this population at passage 2, characterized by a trisomic chromosome 27. For SePOP2-4, the closest pre-existing karyotype is kar4, which has a pentasomic chromosome 31. At passage 2, SePOP4 (brown nodes) is almost identical to kar4, being one step away from this karyotype. This single somy difference is due to a trisomy in chromosome 27 which is not present in kar4. From passage 2 to 3, SePOP4 accumulated 3 extra somy changes (trisomy in chromosomes 6 and 18 and a tetrasomy in chromosome 23). For SePOP2 (green nodes) and SePOP3 (red nodes), the first aneuploidy changes emerged at passage 3 and had already 2 and 3 somy differences compared to kar4 respectively. Altogether, our single-cell data suggest that (i) aneuploidy changes observed in the Sb^III^-exposed populations are explained by the selection of pre-existing aneuploid cells, complemented by additional somy changes generated de novo during the experiment and (ii) that the aneuploid SePOP1-4 would have a polyclonal origin.

### Changes in aneuploidy associated with Sb^III^ selection have a polyclonal origin

In order to document the cell population dynamics during adaptation to the stress generated by Sb^III^, we applied the use of cellular barcodes to track the evolution of hundreds of clonal barcoded lineages during flash selection. In summary, we generated a barcoded promastigote population consisting of 453 different traceable lineages. The frequency of each lineage in each SePOP was monitored by amplicon sequencing. The flash selection induced a fourfold reduction in lineage diversity that stabilized between passages 3 to 4, leaving between 101 to 131 of detectable lineages (fig. S3A).

To distinguish frequency changes associated with Sb^III^ pressure from other sources of variation, such as stochastic loss of lineages due to passaging, we normalized the frequency of each lineage in each SePOP to their respective average frequency in the cPOP at each passage (Fig. 2A). Here we refer to this parameter as “Sb^III^-associated frequency change”. In this sense, lineages that e.g., die or are negatively affected under antimony-pressure but not under standard in vitro conditions display a negative Sb^III^-associated frequency change, while lineages that are eliminated over time in both cPOP and SePOP display similar values. This analysis showed that a large fraction of lineages representing a total 74,6% to 84,8% of the initial population was negatively affected or completely eliminated during Sb^III^ pressure (fig. 2B). This fraction was represented by a total of 362-395 lineages, including 303 lineages consistently affected negatively in all four SePOP (fig. 2C, left panel, red), suggesting that this subset of lineages had a lower tolerance to Sb^III^-generated stress. Interestingly, the fraction of positively affected lineages was relatively small, representing 5,2% to 8,4% of the initial population (Fig. 2B), with only 60 lineages displaying a frequency higher in the drug-exposed group compared to the controls at passage 5. This subset of lineages became dominant to represent 77,7% to 97,4% of the final populations at passage 5. Most of the positively affected lineages were enriched in only one of the SePOPs (Fig. 2C and fig. S3B), suggesting that (i) a subset of lineages was fitter to Sb^III^ prior the drug exposure and (ii) their expansion was stochastically driven.

**Fig. 2.**
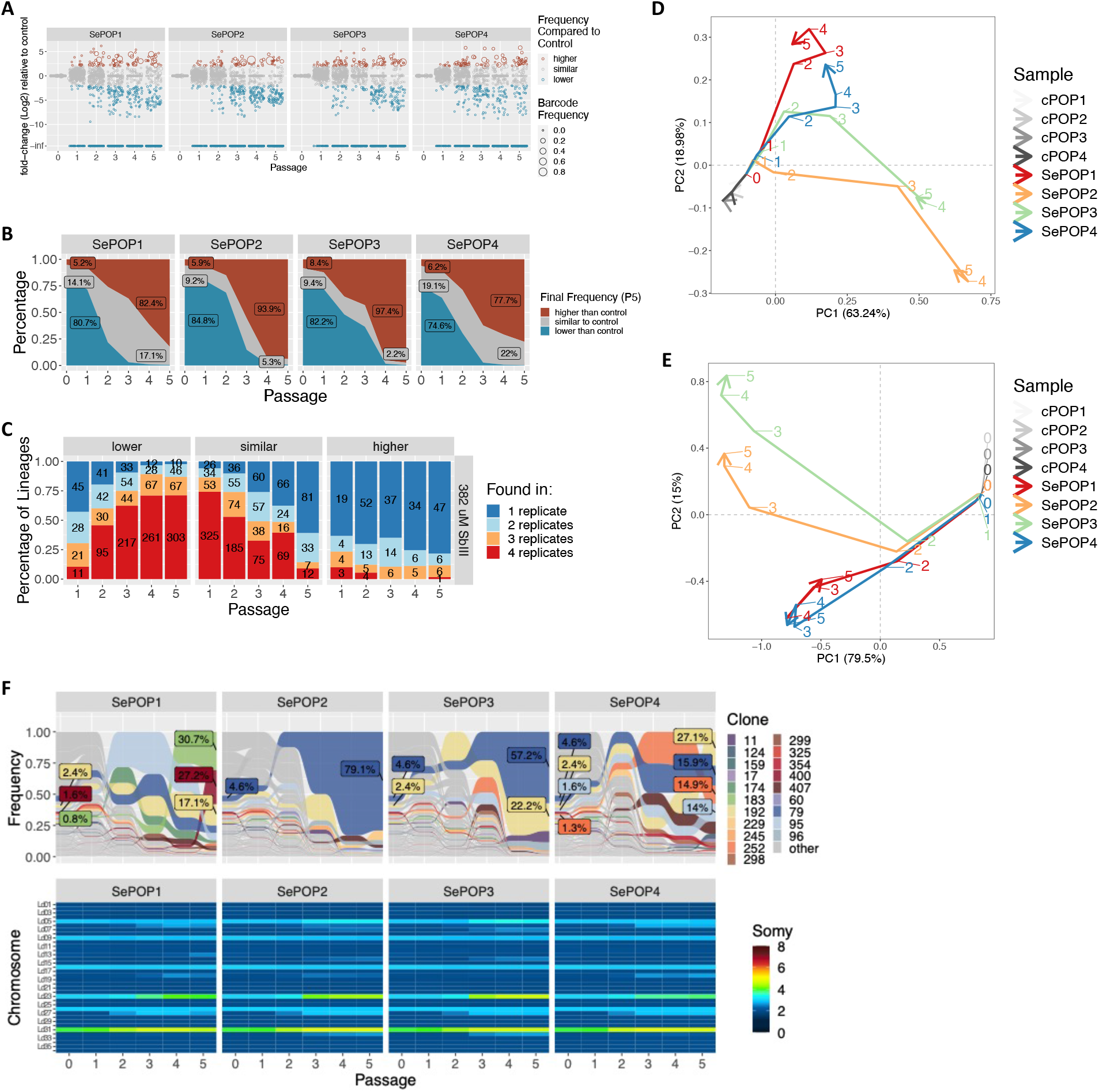
Clonal dynamics of Sb^III^ adaptation revealed with cellular barcodes. **A**. Fold-change of the frequency of each clone in the Sb^III^-exposed group relative to their frequency in the control group in the same timepoint (Sb^III^-associated fold change). Each dot represents a barcoded lineage. Lineages with a log2 Sb^III-^associated fold change smaller than -2 or greater than +2 were considered negatively and positively affected respectively. Lineages with fold-change at -infinite are lineages that were eliminated under drug pressure. **B**. Fraction of lineages that by passage 5 were either positively affected (red) or negatively affected (blue) by Sb^III^ pressure. **C**. Evaluation of the consistency of the Sb^III^-associated fold change scores among replicates. The bars represent the proportion of clones that had a particular Sb^III^-associated fold-change effect (higher, lower or same as in control) in 1, 2, 3 or 4 replicates (dark blue, light blue, orange and red respectively). The numbers in the bars indicate the absolute number of barcodes. **D**. Trajectory principal component analysis (PCA) of the changes in clonal composition over time in each population. The PCA was based on the frequency of each lineage identified in each sample. The numbers indicate the passages at which each sample was collected, while colors indicate the populations. **E**. Trajectory PCA depicting the changes in aneuploidy in all samples. Numbers indicate the passage number at which a sample was collected for WGS. Due to strong similarity between controls, they are not well visible in the PCAs as they cluster very close to each other. **F**. Frequency of each barcoded lineage along the 5 passages under Sb^III^ pressure (top panel). Only barcodes that reached frequencies higher than 1% at passage 5 in at least one of the replicates were colored. A repetition of fig. 1A is included for comparison (bottom panel).

In an attempt to link the evolution of lineages as revealed by barcoding sequencing with the aneuploidy modifications observed in the bulk whole genome sequencing of SePOP1-4, we processed each dataset with a ‘trajectory’ principal component analysis (PCA) and compared them. The former one revealed that lineage composition progressively diverged between replicates, with SePOP1/4 deviating further from SePOP2/3 at later passages (Fig. 2D). Interestingly, this PCA based on lineage composition resembled the aneuploidy-based PCA shown in fig. 2E, with SePOP1/4 and 2/3 clustering separately, suggesting that changes in aneuploidy coincide with changes in lineage composition. This observation is supported when comparing the absolute frequency of the lineages with the aneuploidy changes observed in bulk at each timepoint (fig. 2F and fig. S3C). Here we observe that the major aneuploidy alterations arise at passages 2 to 3, coinciding with the moment where the most dominant lineages in the initial population are depleted while other lineages expand. It is also noticeable that the aneuploidy changes in SePOP2/3 are almost identical, and these two populations were dominated by the same lineage (lineage 79). Conversely, SePOP1/4 also share similar aneuploidy changes, though different from replicates 2 and 3, but lineage composition seems to be less similar.

### Adaptation to miltefosine flash selection is associated with polyclonal selection of pre-existing nucleotide variants

The results described above demonstrated the importance of aneuploidy for parasite adaptation to high Sb^III^ pressure together with the polyclonality of corresponding molecular adaptations. We aimed here to verify if the same features would be observed with another anti-leishmania drug, miltefosine. In contrast to Sb^III^, there was – at least before present study – no pre-adaptation known to miltefosine in the BPK282 strain, which is considered very susceptible to the drug (22).

In order to initiate a flash selection with miltefosine we first determined which was the highest concentration of the drug in which viable parasites could still be recovered. This was done by submitting BPK282 promastigotes to a 1:2 serial dilution of miltefosine ranging from 100 µM to 3,125 µM (2,5 × 10^6^ promastigotes per concentration) in complete culture medium (HOMEM + 20% FBS) in 4 replicates per condition (MIL-exposed populations, MePOP1-4 Fig.S4a). The cultures with 50 µM and 100 µM of miltefosine did not display live parasites even after 1 month post addition of the drug (no passaging). However, in the cultures with 25 µM of MIL, although no live parasites could be detected by microscopy until the 10^th^ day, viable promastigotes started to arise afterwards, and by the 17^th^ day post addition of MIL, MePOP1-4 displayed a cell density comparable to the controls (data not shown). These populations of survivors displayed such a higher tolerance to miltefosine that an attempt to determine their IC_50_ against the drug was not successful as the cultures did not suffer a reduction in viability even in the highest concentration used in the test (75 µM – Fig.S4b). Their respective IC50s were determined at passage 2 by including an additional concentration of 150 µM in the test, and was defined as an average of 81,78 µM in MePOP1-4 and 21,9 µM in the control group (p < 0,001 – Fig. S4C).

When looking for genomic changes after bulk sequencing, we found that the strong bottleneck associated with miltefosine exposure did not lead to any major alteration in aneuploidy, as MePOP1-4 displayed the same profile as the initial population even after 9 passages under miltefosine pressure at 25 µM (Fig.3A). Only after exposure of the population to 100 µM for 4 passages, dosage increases were observed in several chromosomes, with each MePOP displaying a different profile, although they all shared the amplification of chromosome 31 (Fig.3A). In contrast to the Sb^III^ model, relevant SNPs were encountered between MePOP1-4 and the controls, already from the first passage. In particular, in the 4 selected populations, a missense mutation arose in the phospholipid-transporting ATPase1-likeprotein gene, which is also known as the miltefosine transporter (LdMT – ID: LdBPK_131590.1), with MePOP1, 3 and 4 displaying a substitution of a glycine by an aspartate at amino acid (Gly160Asp), while MePOP2 displayed a different mutation in the same gene with an insertion of a stop codon in place of a glutamate in the amino acid 1016 (Glu1016Stop – Fig. 3B).

**Fig. 3.**
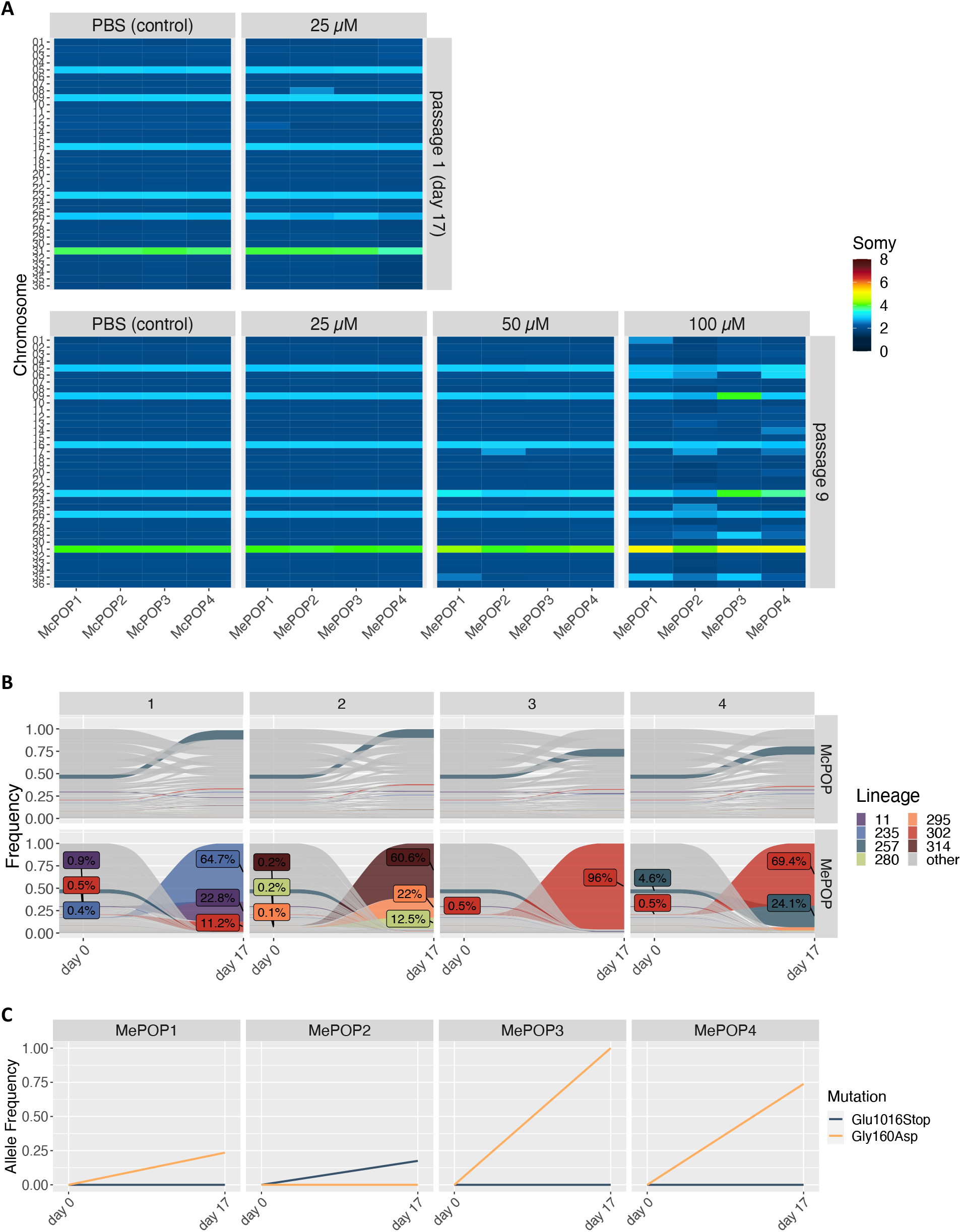
Flash selection with miltefosine. **A**. Bulk aneuploidy profile of the populations kept under different concentrations of miltefosine at passages 1 (upper panel) and passage 9 (bottom panel). **B**. Tracing of lineages before (day 0) and after 17 days (passage 1) under selection with 25 µM miltefosine (MePOP) or with PBS as control (McPOP). Only lineages that reached frequencies higher than 1% in at least one population in the last timepoint (day 17) are colored. Colored labels display the frequency of some lineages at day 0 and day 17. **C**. Allele frequency of the Gly160Asp and the Glu1016Stop mutations identified in the miltefosine-transporter gene (LdMT) in the drug-exposed populations.

As the BPK282 population used in the MIL-flash selection was the same barcoded population as in the Sb^III^-flash selection, we also monitored clonal dynamics between passages 0 (before miltefosine-exposure) and 1 (17 days under miltefosine pressure). This revealed that the bottleneck generated by miltefosine exposure was even stronger than the one associated with Sb^III^ exposure, with only 7 lineages surviving in at least one of the MePOP replicates, with one specific lineage (lineage 302) being present in 3 of the 4 replicates (MePOP1, 3 and 4) at passage 1 (Fig.3C). Interestingly, the frequency of lineage 302 in all 3 replicates where this lineage survived coincided with the allele frequency of the Gly160Asp mutation, suggesting this was a pre-existing mutation present in this lineage. The different mutation seen in MePOP2 also coincides with the fact that this replicate was dominated by different clones.

## Discussion

In the present study, we have applied multiple technologies (i) to understand how environmental stresses, in particular high drug-pressure, promote changes in aneuploidy in vitro, and (ii) to assess the evolutionary dynamics of these aneuploidy changes. Our flash selection models were quite different from conventional drug resistance selection experiments in which parasites are progressively submitted to increasing concentrations of drug (22–24). In our Sb^III^ flash selection model, we consistently observed drastic changes in aneuploidy emerging in a short period of ∼3 weeks, affecting multiple chromosomes, and with different karyotypic outcomes between experimental replicates and repetitions. In almost all cases however, we observed the recurrent dosage increase of a set of chromosomes, including chromosomes 23 and 31. Interestingly, the dosage changes of these chromosomes are commonly reported in several studies in which antimony resistance was selected in vitro through progressive increase of antimony dose, including other *L. donovani* strains (18) as well as other *Leishmania* species such as *L. infantum, L. guyanensis, L. braziliensis and L. panamensis* (25–27). Chromosome 23 bears the MRPA genes which encode an ABC-thiol transporter involved in the sequestration of Sb-thiol conjugates into intracellular vesicles (28). Amplification of MRPA genes through extra-or intra-chromosomal amplification is a well-known driver of experimental Sb^III^ resistance. The line here used (BPK282) is remarkably pre-adapted to Sb^III^ (18) – like other strains of the Gangetic plain – thanks to a pre-existing intra-chromosomal amplification of MRPA genes encountered in 200 sequenced *L. donovani* isolates of that region (13). The recurrent dosage increase of chromosome 23 observed here under Sb^III^ pressure is a rapid way to further amplify the MRPA gene and this mechanism was likely selected by the parasite instead of further amplifying MRPA genes intra-chromosomally. Noteworthy, even in parasites in which MRPA was artificially deleted, chromosome 23 still consistently display increase in copy number in populations selected for antimony-resistance, suggesting that other genes in this chromosome beyond MRPA might be relevant to antimony tolerance (29). Among others, this chromosome also carries the ABCC2 gene, another ABC-transporter which over-expression promotes an increase in Sb^III^ tolerance in parasites in which MRPA was deleted.

The evolutionary dynamics of aneuploidy changes under Sb^III^ pressure was studied with 2 approaches. First, we used high throughput single-cell genome sequencing to evaluate if drug-associated aneuploidy changes were due to the selection of pre-existing cells with karyotypes that already bear those aneuploidies or if they were generated de novo during the experiment. Although we found – prior to Sb^III^ exposure – two single-cells bearing the 2 chromosome amplifications consistently observed in populations under Sb^III^ pressure (i.e., tetrasomy in chromosome 23 and pentasomy in chromosome 31), none of the karyotypes observed in single-cells was identical to those identified under Sb^III^ pressure. We cannot exclude the existence of pre-existing karyotypes present at frequencies lower than the detection limit of the single-cell genomics data (here, 1 in 864 or 0,116%). However, the lineage tracing data indicate that lineages that were selected under Sb^III^ exposure were already at frequencies above this detection limit in the population submitted to single-cell genome sequencing (see fig. 2F, passage 0). Thus, the alternative and most likely explanation is that the 4 (bulk) karyotypes selected under drug pressure indeed originated from pre-existing karyotypes but underwent further and rapid de novo changes in chromosome number under Sb^III^ pressure. This is further supported by the minimum spanning tree displaying the number of somy changes events in karyotypes of single-cells and those of the 4 Sb^III^-exposed cell populations: SePOP1 branched directly from kar16 sharing the same aneuploidy profile with this karyotype at passage 2 while SePOP2-4 branched closely to the pre-existing karyotype kar4, differing only by 2-3 somy changes, which is compatible with the estimated high rate of somy change (0.002-0.0027 changes/generation/cell, (20)). This tree also suggested a polyclonal origin of the 4 karyotypes selected under Sb^III^ pressure, as SePOP1 and SePOP2-4 branched separately on the tree. Secondly, we used clone tracing data based on our new cell barcoding system (see supplementary fig. 1). From 453 different traceable lineages, 303 consistently disappeared during Sb^III^ exposure and 60 showed an increased frequency in at least one replicate. Most of these positively affected lineages were enriched in only one of the SePOP replicates, suggesting (i) higher tolerance to Sb^III^ in a subset of lineages that reproducibly survived the flash selection and (ii) further expansion of these surviving lineages being stochastically driven. Interestingly, changes in clonal composition in each SePOP coincide with the moments where changes in aneuploidy are observed in these populations, suggesting that these aneuploidy changes are due to the emergence of subsets of fitter lineages.

Finally, we assessed the role and dynamics of aneuploidy under strong pressure of another drug, miltefosine. The flash selection performed with miltefosine revealed a contrasting scenario where aneuploidy remained unchanged even after a stronger bottleneck induced by the drug at passage 1, 25 µM and illustrated by the strong decrease in barcode diversity (from 453 to 7 lineages). Aneuploidy changes appeared only at passage 9 (∼45 generations later) and under a miltefosine pressure of 100 µM. This contrasted with the apparently stable aneuploidy profile of the populations observed, also at passage 9, under 25 µM and 50 µM miltefosine pressure. At 100 µM, aneuploidy changes were specific to each of the 4 MePOP replicates, with the exception of chromosome 31 that consistently showed a higher somy than the control. The fact that an increase in copy number of chromosome 31 was observed under strong Sb^III^ and miltefosine pressure, as well as under pressure of other drugs (23) might indicate that the dosage increase in this chromosome has a general role against multiple types of stresses.

Aneuploidy dynamics is thus clearly dependant of the nature and strength of the environmental stress. The contrast here observed between the aneuploidy dynamics under Sb^III^ and miltefosine pressure could be explained by 2 main factors. (i) Aneuploidy changes are not selected under 25-50 µM miltefosine pressure, because another mechanism can promote survival of the parasites. Indeed, two independent mutations were already observed at P1 in LdMT, the main miltefosine transporter gene (phospholipid-transporting ATPase1-likeprotein gene) and this in the 4 replicates. Disruptive mutations in this gene are known to confer resistance to miltefosine in *Leishmania* (30). Interestingly, the Gly160Asp mutation also correlated with the frequency of a specific lineage (lineage 27) and appeared in 3 of the 4 MePOPs, indicating that this was a pre-existing mutation found in that lineage. (ii) The adaptive importance of aneuploidy may also depend on the needed gain or loss of expression for driving drug tolerance. The MRPA is responsible for sequestration of Sb^III^ and thus a higher expression will increase parasite fitness under drug pressure; this should occur by gene dosage and it can easily and rapidly be achieved by multiplying the effect of the MRPA amplification already present in BPK282, by increasing the somy of chromosome 23. In contrast, LdMT is responsible for uptake of miltefosine and in this case, a reduction in expression is driving resistance. In a previous study, this was initiated at low miltefosine pressure by decreasing the somy of chromosome 13 (bearing LdMT), in a strain that was trisomic for that chromosome, from 3 to 2 copies (22). Here chromosome 13 was already disomic and could probably not further decrease in somy and loss of function was achieved via the SNP ‘path’.

In conclusion, we have gathered data showing the importance of aneuploidy changes in the adaptation to strong environmental stress, here high drug pressure in vitro. These changes occurred at different time points of our experimental evolution study, probably depending on the occurrence of mutations increasing parasite fitness and the type of function changes (increase or reduction of expression) needed to drive adaptation. Drug-selected aneuploidy changes showed to have a polyclonal origin as shown by our new method of barcoded lineage tracing and by single-cell genome sequencing. Our data support the role of mosaic aneuploidy in generating multiple pre-adapted karyotypes that can be further modulated de novo during drug exposure, potentially due to a stress-induced increase in chromosome instability as seen in other organisms (31–33). Further research with longitudinal single-cell genome sequencing combined with lineage tracing are needed in order to validate these hypotheses. In addition, our studies should be complemented with in vivo models, where different environmental stresses can be encountered by the parasite and where the biology of the parasite is different (for instance low replication rate or deep-quiescence (34)). Finally, the new barcoding method developed and applied here could be used for characterization of clonal dynamics of *Leishmania* during colonization of in vivo environments, such as sandfly vectors and mammal hosts. It can also provide important information of the role of polyclonality on tissue tropism, as has been recently shown for *Toxoplasma gondii* (35).

## Materials and Methods

### Abbreviations

In the present study, different types of samples are being studied and analysed by bulk or single-cell approaches. To avoid confusion, we list here the main terms used for sample description.

#### SePOP1-4

Sb^III^ exposed Populations; these are populations of cells that are analysed by bulk methods (Whole Genome sequencing, barcode amplification), with 4 replicates. Noteworthy, when we refer to the somy of a given chromosome in a SePOP, this represents an average value calculated on the population of sequenced cells; these values can be integers (most cells of that population show this somy) or intermediate (in that case, there are subpopulations with different somy, for instance a somy value of 2.5 may mean that ∼50% of the population is disomic for that chromosome and ∼50% is trisomic, or other combinations).

#### MePOP1-4

idem but with miltefosine exposure

#### cPOP

control populations not exposed to drugs and maintained in parallel to the drug-exposed populations.

### Promastigote culture

A clonal promastigote population of the *L. donovani* BPK282/0 strain (MHOM/NP/02/BPK282/0 clone 4) was maintained at 26°C in culture with HOMEM medium (Gibco™) supplemented with 20% of heat inactivated foetal calf serum with regular passages being performed every 7 days with 1 in 50 dilutions.

### Flash selection with Sb^III^

A single culture of the barcoded BPK282/0 cl4 strain at 10^6^ promastigotes/ml was divided in 2, where one was further diluted to a final concentration of 5×10^5^/ml with medium containing 764 µM of potassium antimony tartrate (final concentration 382 µM, in the text referred to as Sb^III^) and the other was diluted to the same parasite concentration with a medium containing 0,2% of PBS instead (final PBS concentration 0,1%) as control. These two cultures were subsequently aliquoted into 4 culture flasks each, with a final volume of 5 ml per flask. Each was culture was subcultured every 7 days for a total of 5 passages by transferring 2,5 × 10^6^ promastigotes to a new flask containing fresh medium with either 382 µM Sb^III^ or 0,1% PBS in a final volume of 5 ml. At the end of each passage (day 7), genomic DNA was extracted from ∼10^8^ parasites per flask using the QIAamp DNA Mini Kit (QIAGEN) for subsequent barcode and whole genome sequencing.

### Flash selection with miltefosine

Flash selection with miltefosine was initiated with the same barcoded BPK282 population as in the Sb^III^ flash selection. Flasks containing 5×10^5^ promastigotes per ml (final volume of 5 ml) received miltefosine at 25 µM, 50 µM and 100 µM (4 replicates per concentration) and were maintained without passaging for 1 month, with 4 control replicates kept in parallel. After 17 days, the cultures at 25 µM displayed a full recovery of viability and were maintained for 2 additional passages done every 7 days with 2,5 × 10^6^ promastigotes being transferred to new medium at each passage. At passage 3, the 25 µM cultures were divided in 2, one maintained at 25 µM and the other being exposed to 50 µM of miltefosine. At passage 5, the 50 µM cultures were also divided in 2, this time one kept with 50 µM and another with 100 µM. Cultures were maintained for 4 additional passages until the experiment was stopped.

### Bulk whole genome sequencing

For bulk whole genome sequencing, ∼10^8^ promastigotes were pelleted, washed 3 times with PBS and had their genomic DNA extracted with the QIAamp DNA mini kit (QIAGEN). Sequencing libraries were prepared at GenomeScan (Netherlands) and submitted to 2×150 pair-end sequencing in a NovaSeq™ 6000 sequencer (Illumina). Reads were mapped to the reference LdBPK282-V2 genome (11) and read count was calculated along 5kb bins. Removal of outlier bins and correction of GC-bias were performed as previously established (*36*). Somies were estimated as the median read count per bin of chromosomes normalized by the median read count per bin of the total genome and multiplied by 2.

### Barcode library construction

To generate a library of semi-random barcodes, 20 nmole of an 80-bp single stranded oligonucleotide (5’-ACTGACTGCAGTCTGAGTCTGACAGWSWSWSWSWSWSWSWSWSWSWSWSWSWSWSAGCAGAGCTACGCACTCTATGCTAG-3’) were synthesized by Integrated DNA Technologies™. The barcodes consisted of 15 repeats of A or T (W) – G or C (S) bases and were flanked by conserved regions for barcode amplification (*37*). Second strand synthesis was performed with an extension reaction using the Advantage® 2 PCR Enzyme System with 1 µM (1,2.10^13^ molecules) of the single stranded oligo, 2 mM dNTP mix and 500 nM of primers GD90 and GD92 in a final volume of 50 µL. The sequence information for all primers used in this study can be found in the Supplementary Table 1. The PCR mix was incubated in a thermocycler with denaturation at 95 °C – 1 min, 4 cycles of denaturation 95 °C – 15 s and Annealing/Extension at 68 °C – 1 min, and a final extension for 1 minute at 68 °C. Second strand synthesis was confirmed by gel electrophoresis. This reaction also generated 15-bp sequences at both ends of the barcode-fragment which were necessary for the downstream cloning of the barcode library into a plasmid vector. The vector was a pLEXSY-Hyg2.1 (Jena Bioscience EGE-272) plasmid that was previously modified by inserting an eGFP reporter gene into the NcoI and NotI sites. This vector was digested with NotI and MluI sites and gel purified. The double stranded barcode library was subsequently cloned into the linearized vector using the In-Fusion® HD Cloning Kit (Clontech® Laboratories) following instructions of the manufacturer. The estimated vector:insert ratio for the reaction was 7,25.10^10^:1,8.10^12^ (≈ 1:25) molecules, which was optimized to keep the diversity of the barcode library. The whole cloning reaction was used to transform Stellar™ chemically competent cells (Clontech® Laboratories) using the heat shock method. After transformation, bacteria were seeded in multiple LB-agar plates (⌀ 150 mm) containing 100 μg/ml ampicillin and were grown at 37°C for 16h. All grown colonies were pooled by adding liquid LB medium to the plates and scraping off the colonies with a cell spreader. From this collected bacterial culture plasmids were extracted using the PureLink™ HiPure Plasmid Midiprep Kit (Thermo Fisher Scientific).

### Generation and characterization of the barcoded *L. donovani* population

The barcoded plasmid pool purified from the bacterial cells was further linearized by a SwaI digestion, whereafter the resulting DNA fragments were separated by gel electrophoresis and the required 5821 bp band was purified using the Wizard® SV Gel and PCR Clean-Up System (Promega). In this linear form, this vector is flanked by homology sequences that promotes its genomic integration into the 18S rDNA locus through spontaneous homologous recombination. For cellular barcoding, a *Leishmania donovani* clonal population (BPK282/0 cl4) derived from the MHOM/NP/02/BPK282/0 strain was maintained as promastigotes at 26°C in HOMEM medium (Gibco, ThermoFisher) supplemented with 20% Foetal Bovine Serum, with regular passages done every 7 days at 1/50 dilutions. At passage 30 after cloning, 8×10^7^ promastigotes were transfected with 1 µg (≈ 1,68.10^11^ molecules) of the linearized barcoding library using the Basic Parasite Nucleofector™ kit 1 (Lonza) with the U-033 program. After 2 days post transfection, barcoded parasites were continuously selected with Hygromycin B at a final concentration of 25 µg/ml, and the percentage of cells bearing the barcode vector was monitored by detection of the eGFP expression with confocal microscopy and/or flow cytometry. After two passages (14 days) under hygromycin selection, 22 subclones were derived using fluorescence activated cell sorting (FACS) and submitted to a PCR with primers GD124 and GD125 to amplify the barcode region for a subsequent Sanger sequencing with the primer GD126 in order to determine the presence or absence of a barcode in individual promastigotes as well as to detect potential multiple insertions.

### Barcodes amplification and sequencing

Barcodes were amplified in a two-step PCR approach using the Kapa HiFi HotStart ReadyMix PCR kit (Kapa Biosciences). The first PCR was done with primers NtBCampF and NtBCampR at a final concentration of 10 µM each in a final volume of 100 µL per reaction containing ∼400 ng of the template DNA. This PCR was carried out with a initial denaturation at 95°C for 3 minutes, followed by 18 cycles of denaturation at 98°C for 20 seconds, annealing at 65°C for 30 seconds, and extension at 72°C for 20 seconds, with a final extension at 72°C for 2 minutes. The PCR reactions were purified using 1,45x AMPure XP Beads (Beckman Coulter), eluted in 10 µL of water and directly transferred to a second PCR reaction containing the primers NtIndex-F and NtIndex-R at 10 µM each in a final volume of 50 µL. These primers contain the adapters and index sequences needed for Illumina sequencing. The temperature cycle was 95°C for 3 minutes, followed by 17 cycles of 98°C for 20 seconds, 65°C for 30 seconds, and 72°C for 20 seconds, with a final extension at 72°C for 2 minutes. The final PCR product was purified once more with 1,45x AMPure XP beads and eluted in 20 µL of the Buffer EB (QIAGEN). The libraries were quantified using the KAPA Library quantification kit (Kapa Biosciences) and were pooled at equimolar ratios in a final concentration of 3nM. The library pool was sequenced using a Illumina™ NextSeq 550 sequencer with 2 × 75pb reads targeting on average 5 milion pair-end reads per sample.

### Barcode counting

Sequencing read pairs in which at least one of the reads had an average quality score lower than 20 were removed from downstream analysis. Then, read pairs were merged to form consensus sequences using the FASTP pipeline (*38*). The Bartender pipeline (*39*) was used to generate the barcode count table, using the pattern “GACAG[29-39]AGCAG”, and allowing for 2 mismatches. Downstream analyses were done in R. To distinguish barcodes originating from PCR or sequence errors from true barcodes, we submitted one sample (BPK282 population prior to drug exposure) to 3 independent library preps. After confirming that barcodes appearing in only 1 or 2 libraries were always detected at low frequencies, we considered as true barcodes only those found in all 3 libraries.

### Single-cell genome sequencing

Single-cell genome sequencing was performed using the single-cell CNV™ solution from 10X Genomics. Execution of the protocol and bioinformatic analysis were done as previously described (20).

### Dose response assay against antimony and miltefosine

Dose-response assays were performed as previously described (40). In summary, logarithmic-stage promastigotes were seeded in 96-well plates at final concentration of 10^5^ parasites per well in supplemented HOMEM medium per well. Parasites were exposed to different concentrations of miltefosine ranging from 3 µM to 75 µM (first attempt) or 150 µM (second attempt). An extra row of wells was kept with 0,2% PBS instead of antimony as a control. Parasites were exposed to the drug for 3 days at 26 °C. Afterwards, each plate received 20 µg/ml of resazurin and was further incubated for 4h at 26°C. Then, light absorption was measured at 560nm/590nm excitation/emission Using a Victor X2 luminometer. Assays were independently performed 3 times, with 3 identical plates in each execution serving as technical replicates. When possible, half maximal inhibitory concentrations (IC50) were calculated with GraphPad Prism using a sigmoidal dose-response model with variable slope.

## Author contributions

Conceptualization: GHN, GM, FVB, JCD, MAD

Methodology: GHN, GM, MAD

Investigation: GHN, RG, DVG, IM

Visualization: GHN, PM

Formal analysis: GHN, PM

Supervision: JCD, MAD

Writing – original draft: GHN, JCD, MAD

Writing – review & editing: GHN, PM, GM, FVB, JCD, MAD

## Data and material availability

Custom scripts used for analysis of barcode and whole genome sequencing data are available at https://github.com/gabrielnegreira/LeishBarSeqAndAneuploidy. Scripts for single-cell data are available at https://github.com/gabrielnegreira/scgs-somy. All data are available in the main text or the supplementary materials.

