## Supplementary Material for "The role of aneuploidy and polyclonality in the adaptation of the Protozoan parasite *Leishmania* to high drug pressure"

Gabriel H. Negreira *et al.*

**This PDF file includes:**

Figs. S1 to S4

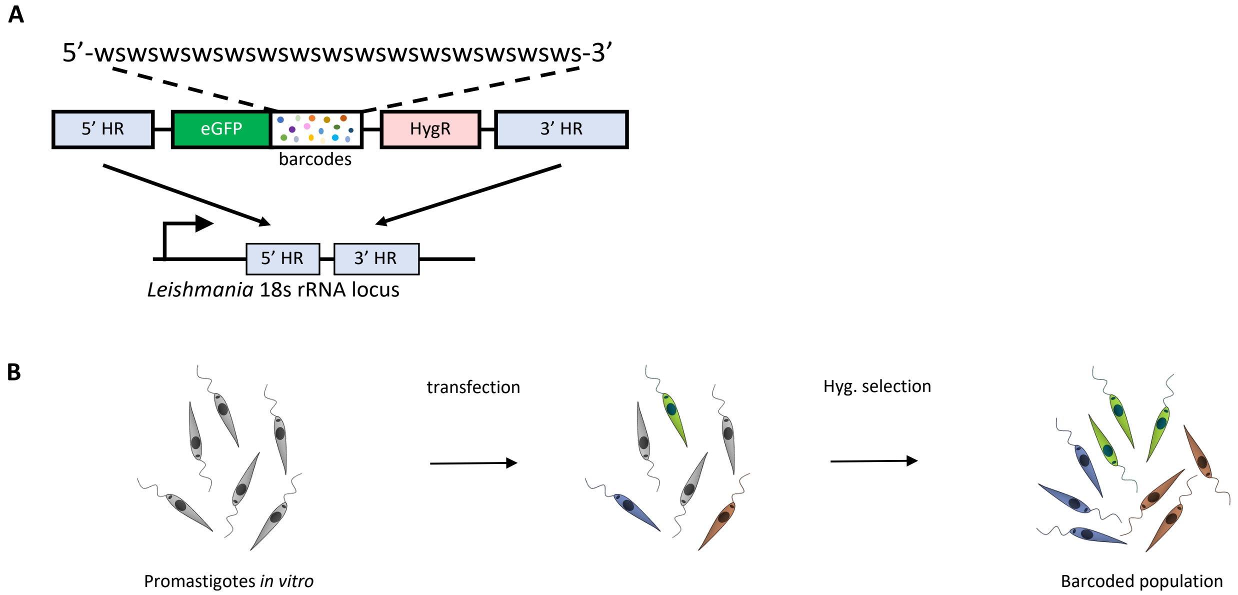


**Fig. S1 - Schematic representing the cellular barcoding strategy. A.** Briefly, a double-stranded oligonucleotide is synthesized bearing a semi-random sequence formed by the alternation of weak (W = A or T) and strong bases (S = G or C) flanked by two fixed sequences that serve as primer binding sites for PCR amplification. This pool of DNA molecules is cloned in a vector that has homology sequences which promote the integration of the vector into the 18s rRNA locus in *Leishmania* genome. **B**. After transfection, barcoded parasites are selected with hygromycin, as the barcoding vector also has a hygromycin-resistance gene. The eGFP gene allow to monitor by flow cytometry the potential presence of remaining non-barcoded cells after hygromycin selection.

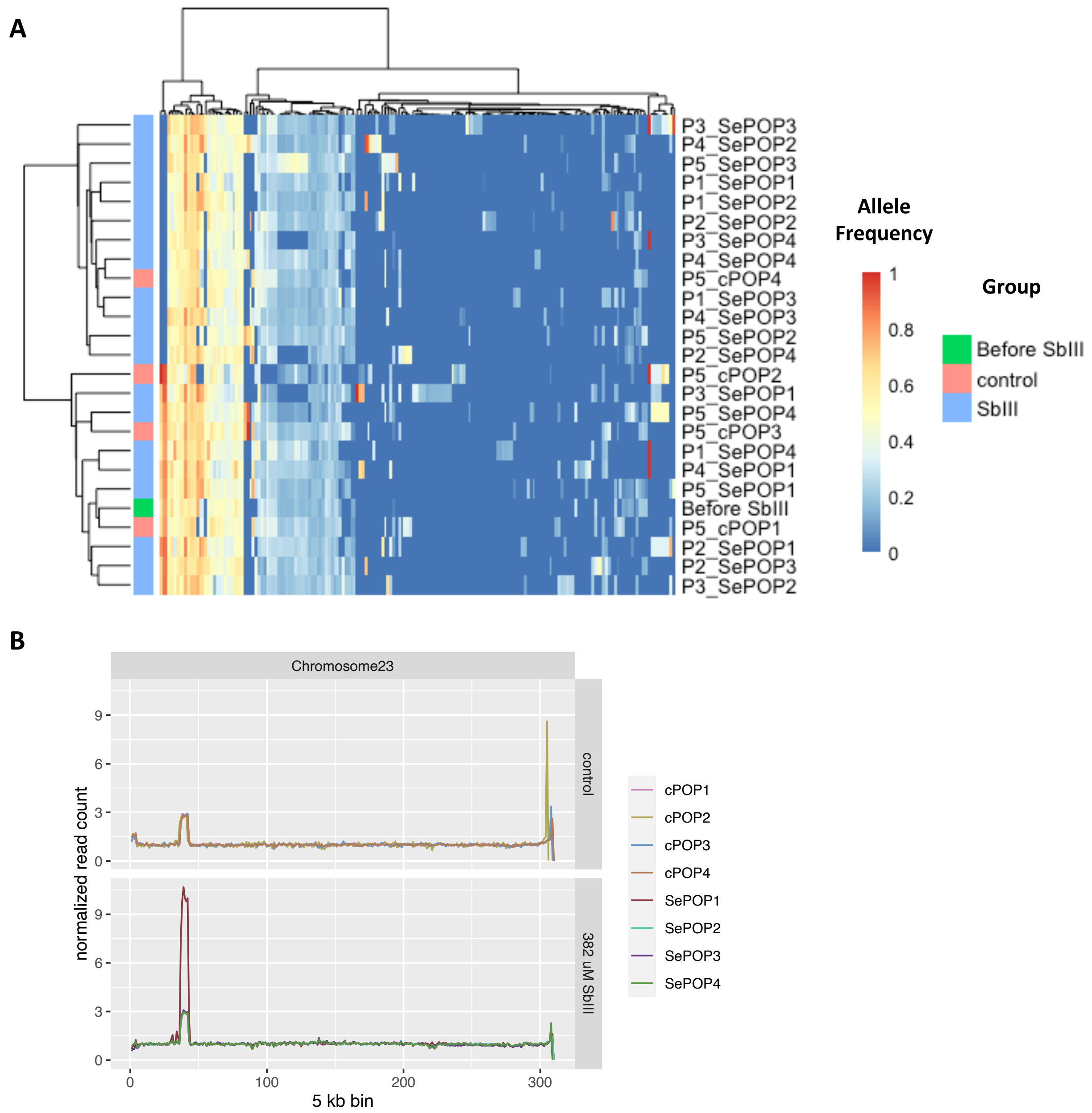


**Fig. S2 – Additional genomic changes associated with the Sb$^{III}$-Flash selection performed on the barcoded BPK282 population. A.** Heatmap depicting the allele frequencies of SNPs and indels identified in protein coding regions compared to the reference genome. An additional annotation bar display to which group (control or Sb$^{III}$-exposed) a sample belongs. Samples are named as Px_cPOPy for controls, and Px_SePOPy for the Sb$^{III}$-exposed groups, with x being the number of passages and y being the replicate number. The initial population is named as 'Before Drug'. **B.** Copy number variation in the MRPA locus. Y axis represent median read count of 5 kb bins in normalized by the median count of chromosome 23 and reflect the average copy number per haploid genome of the MRPA locus.

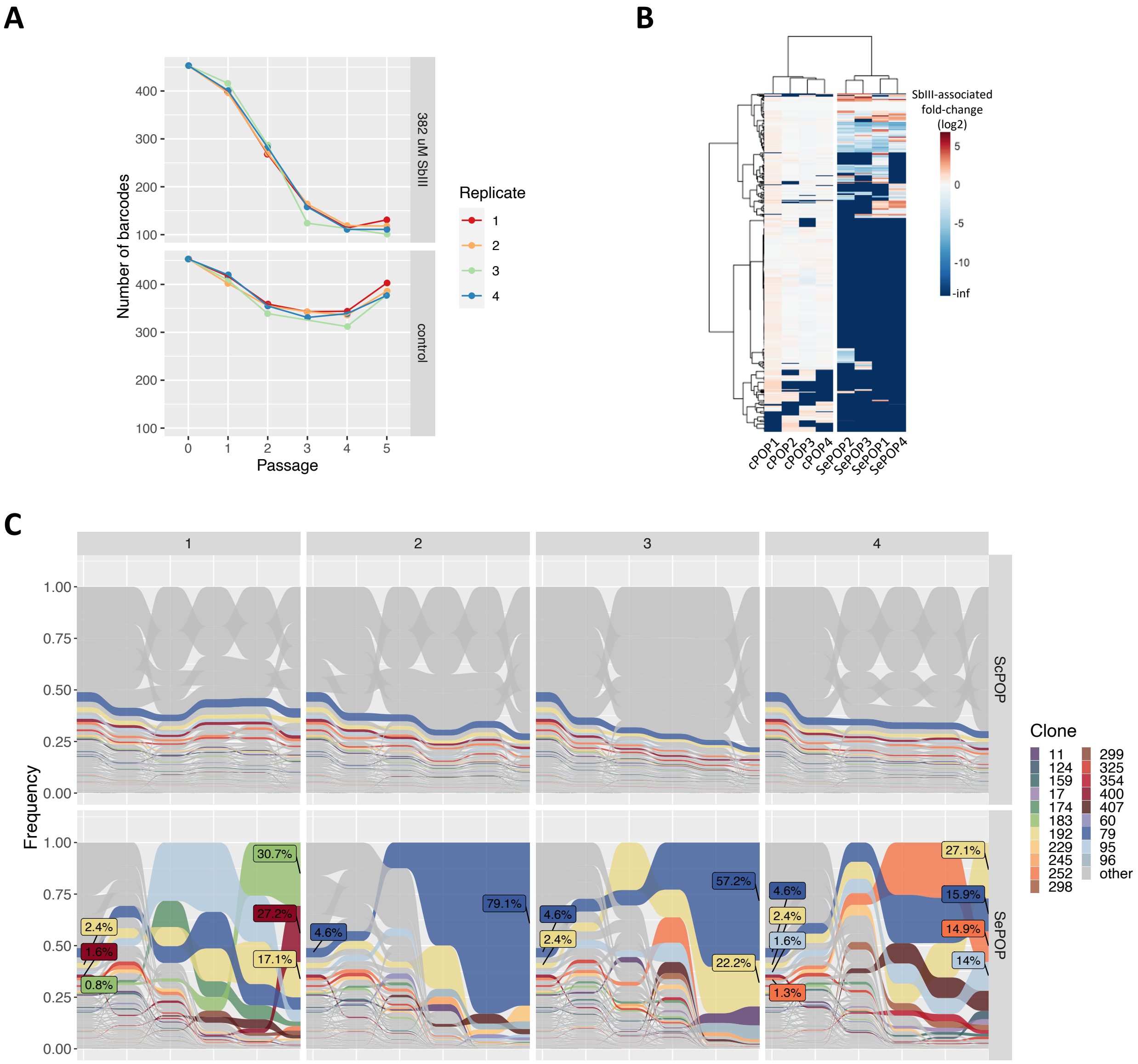


**Fig. S3 – Supporting figures for Fig. 2 of the main text. A.** Total number of different barcodes identified in each population at each timepoint in the $Sb^{III}$-exposed populations (top) and the controls (bottom). **B.** Heatmap displaying the $Sb^{III}$-associated fold-change of each lineage (rows) in each population (columns) at passage 5. **C.** Frequency of each barcoded lineage along the 5 passages in the ScPOP1-4 (top) and SePOP1-4 (bottom) populations. This is similar to main Fig. 2F but include the ScPOP1-4 for comparison. Only lineages that reached a frequency higher than 1% at passage 5 in in at least one of the SePOP populations are colored.

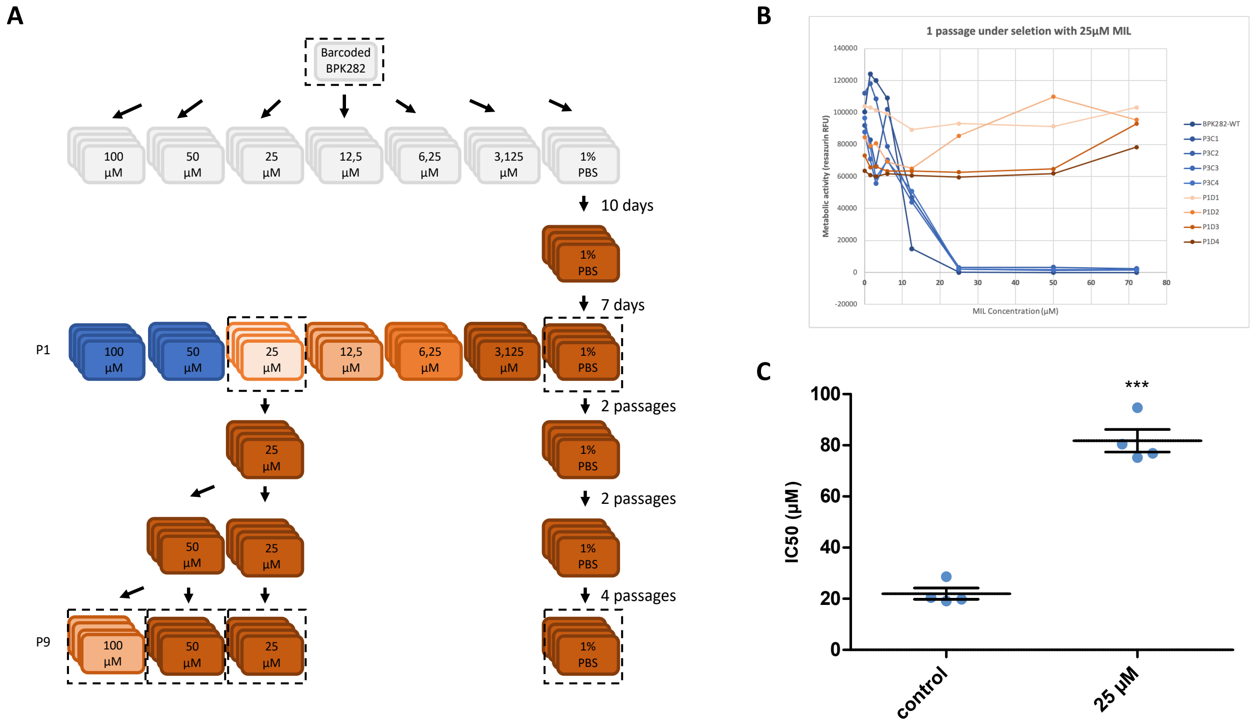


**Figure 4 – Supporting figures for flash selection with miltefosine. A.** Schematic representing the experimental design. Colors indicate the lack of viable parasites (blue) or the relative observed number of parasites at day 7 compared to the controls (darker brown = more cells). Samples submitted to whole genome sequencing are highlighted with a dashed rectangle. **B.** Dose-response to miltefosine of the metabolic activity of the populations at the first passage after exposure to the drug, estimated with the resazurin assay. **C.** Difference in IC50 of the same populations one passage later. *** $p < 0,001$ (T-test).
